# Muscle overexpression of Klf15 via an AAV8-Spc5-12 construct does not provide benefits in spinal muscular atrophy mice

**DOI:** 10.1101/717785

**Authors:** Nina Ahlskog, Daniel Hayler, Anja Krueger, Sabrina Kubinski, Peter Claus, Rafael J Yáñez-Muñoz, Melissa Bowerman

**Affiliations:** Department of Physiology, Anatomy and Genetics, University of Oxford, Oxford, UK; Department of Paediatrics, University of Oxford, Oxford, UK; AGCTlab.org, Centre of Gene and Cell Therapy, Centre for Biological Sciences, Department of Biological Sciences, Royal Holloway, University of London, Egham Hill, Egham, Surrey, UK; Institute of Neuroanatomy and Cell Biology, Hannover Medical School, Hannover, Germany and; Center of Systems Neuroscience, Hannover, Germany; School of Medicine, Keele University, Staffordshire, UK; Institute for Science and Technology in Medicine, Stoke-on-Trent, UK and; Wolfson Centre for Inherited Neuromuscular Disease, RJAH Orthopaedic Hospital, Oswestry, UK

## Abstract

Spinal muscular atrophy (SMA) is a neuromuscular disease caused by loss of the survival motor neuron (*SMN*) gene. While there are currently two approved gene-based therapies for SMA, availability, high cost, and differences in patient response indicate that alternative treatment options are needed. Optimal therapeutic strategies will likely be a combination of SMN-dependent and -independent treatments aimed at alleviating symptoms in the central nervous system and peripheral muscles. Krüppel-like factor 15 (KLF15) is a transcription factor that regulates key metabolic and ergogenic pathways in muscle. We have recently reported significant downregulation of *Klf15* in muscle of pre-symptomatic SMA mice. Importantly, perinatal upregulation of *Klf15* via transgenic and pharmacological methods resulted in improved disease phenotypes in SMA mice, including weight and survival. In the current study, we designed an adeno-associated virus serotype 8 (AAV8) vector to overexpress a codon-optimised *Klf15* cDNA under the muscle-specific Spc5-12 promoter (AAV8-*Klf15*). Administration of AAV8-*Klf15* to severe Taiwanese *Smn*^*−/−*^;*SMN2* or intermediate *Smn*^*2B/−*^ SMA mice significantly increased *Klf15* expression in muscle. We also observed significant activity of the AAV8-*Klf15* vector in liver and heart. AAV8-mediated *Klf15* overexpression moderately improved survival in the *Smn*^*2B/−*^ model but not in the Taiwanese mice. An inability to specifically induce Klf15 expression at physiological levels in a time- and tissue-dependent manner may have contributed to this limited efficacy. Thus, our work demonstrates that an AAV8-Spc5-12 vector induces high gene expression as early as P2 in several tissues including muscle, heart and liver, but highlights the challenges of achieving meaningful vector-mediated transgene expression of Klf15.

## INTRODUCTION

Spinal muscular atrophy (SMA) is a devastating childhood neuromuscular disease that leads to early death in the most severe cases ^1,2^. As an autosomal recessive disease, SMA is caused by loss of the *survival motor neuron 1* (*SMN1*) gene due to either mutations or deletions ^3^. While a total deficiency in the SMN protein is embryonic lethal ^4^, humans have a duplicated copy of *SMN1*, termed *SMN2* ^3^, which allows for survival in the absence of the former. However, *SMN2* contains a key C to T transition in exon 7 that leads to its excision in approximately 90% of the transcripts produced, generating a non-functional SMNΔ7 protein that is rapidly degraded ^5,6^. Importantly, the 10% of fully functional full length SMN protein produced from *SMN2* is sufficient to allow survival, albeit not sufficient to prevent neuromuscular degeneration ^7^.

The first genetic therapy for SMA, nusinersen/Spinraza™, was approved in December 2016 by the Food and Drug Administration (FDA) and in June 2017 by the European Medicines Agency ^8^. This antisense oligonucleotide is delivered directly to the central nervous system (CNS) via a lumbar puncture and is aimed at promoting *SMN2* exon 7 inclusion ^9^. Zolgensma® is a single, systemic of delivery of *SMN1* via an adeno-associated virus serotype 9 (AAV9) gene therapy that received FDA approval in May 2019 ^10^. Additional SMN-enhancing pharmacological compounds are also in the pipeline and anticipated to be approved for patient use in the near future ^11^. While the benefits of these SMN-dependent drugs are undeniably remarkable, it is appreciated that they unfortunately do not represent a cure and will have to be supported by additional non-CNS and -SMN therapeutic interventions to provide optimal care to all SMA patients ^12–15^.

SMN-depleted skeletal muscle displays both cell-autonomous and non-autonomous defects ^16,17^ and is therefore an important therapeutic target for SMA. We have recently demonstrated the dysregulated expression of the transcription factor Krüppel-like factor 15 (*Klf15*) in skeletal muscle of SMA mice during disease progression ^18^. KLF15 is crucial in the regulation of skeletal muscle metabolism and ergogenic properties ^19–22^. Specifically, we observed a significant downregulation of *Klf15* expression in pre-symptomatic SMA mice and found that its neonatal upregulation via pharmacological (prednisolone) or transgenic (muscle-specific *Klf15*-overexpression) interventions significantly improved several disease phenotypes in SMA mice ^18^. However, prednisolone has pleiotropic activities and constitutive embryonic overexpression of *Klf15* in skeletal muscle of SMA may have resulted in compensatory mechanisms ^23^. In this study, we thus set out to overexpress *Klf15* in skeletal muscle of neonatal SMA mice via a self-complementary adeno-associated virus serotype 2/8 and the Spc5-12 promoter. While this strategy led to substantial *Klf15* expression in skeletal muscle of SMA mice and control littermates, there were no associated significant improvements in disease phenotypes. Nevertheless, AAV8-*Klf15* injections resulted in pronounced expression as early as post-natal day 2 in several tissues including muscle, liver and heart, highlighting the potential of this specific viral construct for efficient perinatal delivery.

## MATERIALS AND METHODS

### Animals

Wild-type (WT) FVB/N mice were used for initial expression screening. The Taiwanese *Smn*^*−/−*^;*SMN2* (FVB/N background, FVB.Cg-Smn1tm1HungTg(SMN2)2Hung/J, RRID: J:59313) ^24^ and the *Smn*^*2B/−*^ ^25^ mice (generously provided by Dr Lyndsay M Murray, University of Edinburgh) were housed in individual ventilated cages (fed *ad libitum*, 12 hr light: 12 hr dark cycle) at the Biomedical Sciences Unit, University of Oxford, according to procedures authorized by the UK Home Office (Animal Scientific Procedures Act 1986). The viral constructs were diluted in sterile 0.9% saline and administered at the indicated dose at postnatal day (P) 0 by a facial vein intravenous injection ^26^. Litters were randomly assigned to treatment at birth. For survival studies, animals were weighed daily and culled at indicated time points or upon reaching their defined humane endpoint as set out by the Home Office Project Licence. For the *Smn*^*2B/−*^ mice, weaned mice were given daily wet chow at the bottom of the cage to ensure proper access to food. Sample sizes were determined based on similar studies with SMA mice.

### Sc-AAV2/8-Spc5-12 constructs

The generation of the self-complementary adeno-associated virus serotype 2/8 (scAAV2/8) vectors and quality control were performed by Atlantic Gene Therapies (Nantes, France). The synthetic Spc5-12 promoter ^27^ was used to drive the expression of eGFP or a codon-optimised *Klf15* sequence: agcgcttcaccacaggctcggccaggccagcatggttgatcatctgctgcctgtggacgagacattcagcagccctaagtgttctgtgggctacctgggcgacagactggcctctagacagccttaccacatgctgccctctccaatcagcgaggacgactccgatgtgtctagcccttgtagctgtgcctctcctgacagccaggccttctgtagctgttactctgctggacctggacctgaggctcagggctctatcctggatttcctgctgagcagagctacactcggctctggcggaggatctggcggaatcggagattcttctggccctgtgacatggggctcttggagaagggctagcgtgcccgtgaaagaggaacacttctgcttccctgagttcctgagcggcgacaccgatgacgtgtccagacctttccagcctacactggaagagatcgaagagttcctcgaagagaacatggaagccgaagtgaaagaagcccctgagaacggctcccgcgacctggaaacatgttctcagctgtctgccggctctcacagaagccatctgcaccctgaaagcgccggcagagagagatgtacacctcctccaggtggaacatctggcggcggagcacaatctgctggcgaaggacctgctcatgatggacctgtgcctgtgctgctgcaaatccagcctgtggctgtgaagcaagaggctggaacaggaccagcttctcctggacaggctcctgaatctgtgaaggtggcccagctgctggtcaacatccagggacaaacattcgccctgctgcctcaggtggtgcccagcagtaatctgaacctgcctagcaagttcgtgcggatcgctcctgtgccaatcgctgctaagcctatcggatctggctctcttggtcctggaccagctggactgctcgtgggacagaagttccctaagaaccctgccgccgagctgctgaagatgcacaagtgtacattccccggctgctccaagatgtataccaagtcctctcacctgaaggcccacctgagaaggcataccggcgagaagcctttcgcttgcacatggcctggatgtggctggcggttcagcagatctgatgagctgagcaggcaccgcagatctcacagcggagtgaagccataccagtgtcctgtgtgcgagaagaagttcgccagaagcgaccacctgtccaagcacatcaaggtgcacagattccctagaagcagcagagccgtgcgggccatcaattgactgcagaagctt.

### C2C12 cell line

C2C12 myoblast cells ^28^ were maintained in growth media consisting of Dulbecco’s Modified Eagle’s Media (DMEM) supplemented with 10% fetal bovine serum and 1% Penicillin/Streptomycin (all Life Technologies). For AAV transduction experiments, growth media was changed to differentiation media consisting of DMEM, 2% horse serum, and 1% Penicillin/Streptomycin (all Life Technologies). Cells were allowed to differentiate for 3 days, after which they were transduced with a MOI of 1E10^5^. Cells were harvested 3 days post-transduction for molecular analyses (flow cytometry or qPCR, as described below).

### qPCR

Quadriceps muscles, liver, and heart were harvested at the indicated time points during disease progression and immediately flash frozen. RNA was extracted with the RNeasy MiniKit (Qiagen). For C2C12 cells, the media was removed and cells were washed with PBS before being directly lysed as per instructions within the RNeasy MiniKit (Qiagen). Reverse transcription was performed using the High-Capacity cDNA Reverse Transcription Kit (ThermoFisher Scientific). qPCR was performed using SYBR green Mastermix (ThermoFisher Scientific) and primers for the codon-optimised *Klf15* sequence (Forward: AGGACCTGCTCATGATGGAC; Reverse: TGTTTGTCCCTGGATGTTGA). *RNA polymerase II polypeptide J* (*PolJ*), was used as a validated stably expressed housekeeping gene ^29^ (Forward: ACCACACTCTGGGGAACATC; Reverse: CTCGCTGATGAGGTCTGTGA). All primers were ordered from Integrated DNA Technologies.

### Flow cytometry

Differentiated C2C12 cells 3 days post-transduction (AAV8-GFP, MOI of 1E10^5^) and untreated cells were trypsinized and washed in fluorescence-activated cell sorting (FACS) buffer (phosphate buffered saline (PBS) supplemented with 2% bovine serum albumin (BSA) and 0.05% sodium azide). Cells were pelleted by centrifugation at 300 x g for 5 minutes. The final cell pellet was resuspended in 200 μ of FACS buffer and the cell suspension was further diluted 1:2 in FACS buffer before detection in the Cytek DxP8 flow cytometer (Cytek® Biosciences). Cell viability was tested by adding 1 µl Sytox Red (Thermofisher) to the final cell suspension for 5 minutes at RT. The gating strategy included gating around the Sytox Red (RedFL1 channel) negative population following FCS/FCSW doublet exclusion. The remaining population was assessed for a GFP-shift by recording in the BluFL1 channel. A total of 10,000 cells was recorded for each sample replicate. Data was analysed using Flowjo 10 software (TreeStar Inc.).

### Immunocytochemistry and immunohistochemistry

C2C12 cells transduced with the *eGFP*-expressing AAV vector were imaged live with a DM IRB microscope (Leica).

Quadriceps muscles were harvested at the indicated time points during disease progression, fixed in 4% paraformaldehyde, cryopreserved in 30% sucrose, and cryosectioned at a thickness of 12 μM. The sections were immunostained with chicken anti-GFP antibody (1:3000, Abcam) and detected with Alexa-488-conjugated anti-chicken secondary antibody (1:5000, Life technologies). Images were taken with an Olympus Fluoview FV1000 confocal microscope and processed with Fiji ^30^.

### Statistics

All statistical analyses were performed using GraphPad Prism version 8.1.1 software. When appropriate, a Student’s unpaired two-tailed *t test* or a two-way ANOVA followed by an uncorrected Fisher’s LSD multiple comparison test was used. Outliers were identified via the Grubbs’ test and subsequently removed. Instances of outlier removal are detailed in the relevant figure legends. For the Kaplan-Meier survival analysis, a log-rank test was used.

## RESULTS

### Muscle-specific *Klf15* expression with a scAAV2/8-Spc5-12 viral vector

To specifically induce *Klf15* expression in skeletal muscle, we utilized a self-complementary adeno associated virus serotype 2/8 driven by the synthetic muscle-specific promoter Spc5-12 ^27^ (scAAV2/8-Spc5-12-*Klf15*, henceforth termed AAV8-*Klf15*). This combination of AAV and promoter has previously successfully been used for gene delivery to muscles for treatment of the muscle disorder Duchenne muscular dystrophy (DMD) ^31^. A control scAAV2/8-Spc5-12-*GFP* construct was also generated (henceforth termed AAV8-*GFP*).

We first examined the transduction ability of the AAV8-*GFP* construct in differentiated C2C12 myoblasts ^28^. The cells were transduced with AAV8-*GFP* (multiplicity of infection (MOI) 1E10^5^) for 3 days and assessed for GFP expression compared to untreated cells. Both flow cytometry and live imaging analyses confirm the abundant presence of GFP in AAV8-transduced cells (Fig 1a,b). Differentiated C2C12s transduced with AAV8-*Klf15* (MOI 1E10^5^) demonstrate a significant increased expression of *Klf15* mRNA compared to untreated cells (Fig 1c).

**Fig 1.**
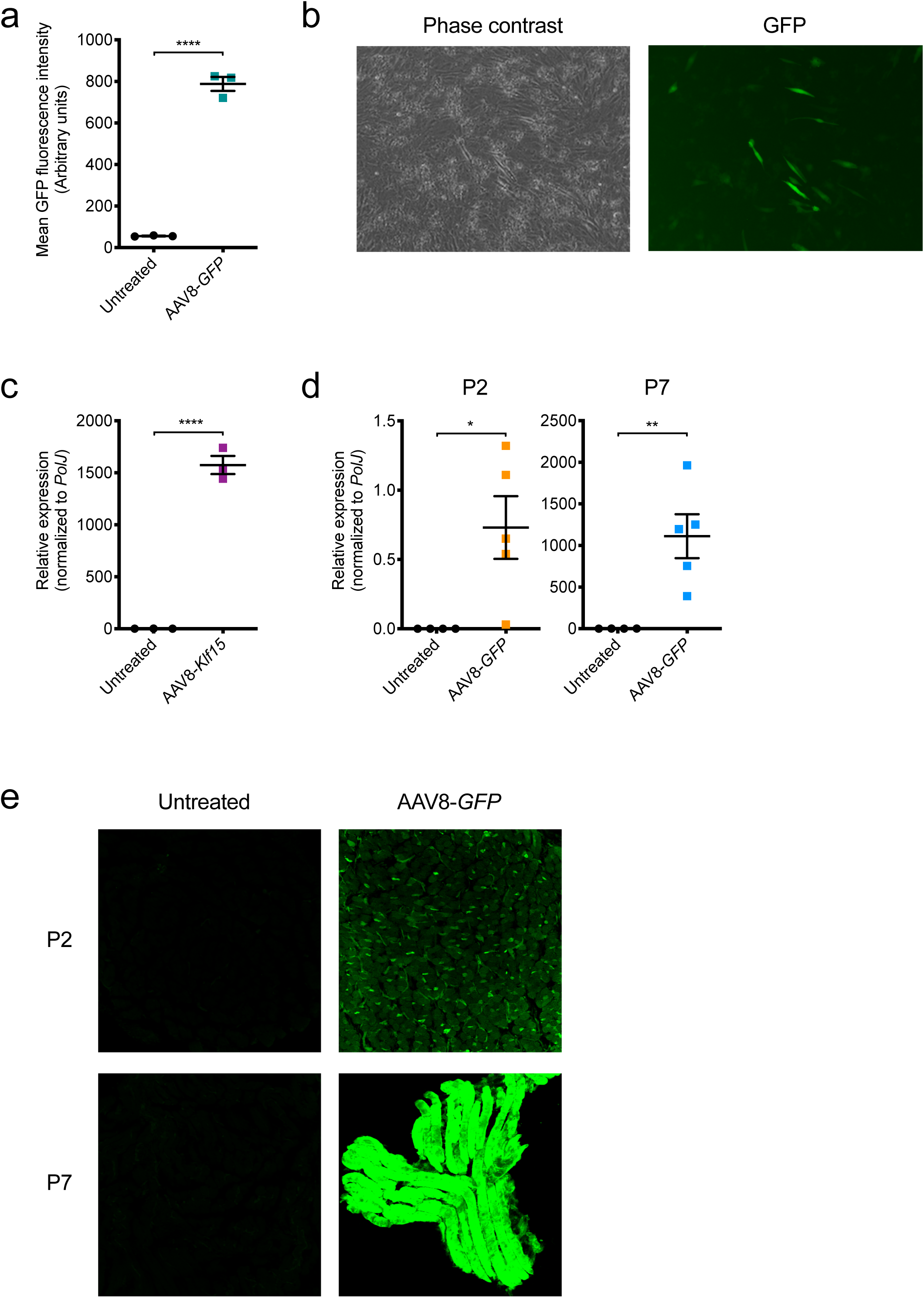
Muscle-enhanced expression with the scAAV2/8-Spc5-12-*Klf15* and-*GFP* (AAV8-*Klf15* and AAV8-*GFP*) viral vectors. **a.** Mean GFP fluorescence intensity (arbitrary units) determined by flow cytometry analysis in differentiated C2C12s, untreated or transduced with AAV8-*GFP* (MOI 1E10^5^) for 3 days. Data are scatter plot and mean ± SEM, n = 3 wells per experimental group, unpaired t test, \*\*\*\**p*<0.0001. **b.** Representative images (phase contrast and GFP fluorescent signal) of differentiated C2C12 cells 3 days post-transduction with AAV8-GFP (MOI 1E10^5^). **c.** qPCR analysis of *Klf15* mRNA expression in differentiated C2C12s 3 days post-transduction with AAV8-*Klf15* (MOI 1E10^5^) compared to untreated cells. Data are shown as scatter plot and mean ± SEM, n = 3 wells per experimental group, unpaired t test, \*\*\*\**p*<0.0001. **d.** qPCR analysis of *GFP* mRNA expression in quadriceps muscles of post-natal day (P) 2 and P7 wild type (WT) animals that received a facial intravenous injection of AAV8-*GFP* (1E11 vg/pup) at P0 compared to untreated WT mice. Data are scatter plot and mean ± SEM, n = 4–5 animals per experimental group, unpaired t test, \**p* = 0.028 (P2), \*\**p* = 0.0076 (P7). **e.** Representative images of cross-sections (P2) and longitudinal sections (P7) of quadriceps immunostained for GFP from P2 and P7 untreated WT animals and WT mice that received a facial intravenous injection of AAV8-*GFP* (1E11 vg/pup) at P0.

To determine if our constructs would also be active at early time points in muscle of neonatal mice, we administered AAV8-*GFP* (1E11 vg/pup) to P0 wild type (WT) pups via a facial vein intravenous injection ^26^. Quadriceps muscles were harvested from injected and non-injected WT littermates at P2 and P7. P2 represents the pre-symptomatic age at which we have observed a significant downregulation of *Klf15* in the Taiwanese *Smn*^*−/−*^;*SMN2* SMA mice ^18,24^ while P7 is considered a late symptomatic time point. qPCR analysis shows a significant upregulation of *GFP* expression at both P2 and P7, albeit variable between animals, with increased levels at the later time point (Fig 1d). Immunohistochemistry of P2 and P7 quadriceps also reveals a time-dependent increased expression of GFP in AAV8-treated animals compared to untreated littermates (Fig 1e). Combined, our experiments in C2C12s and WT mice demonstrate the ability of our viral vectors to induce *Klf15* and *GFP* expression in differentiated skeletal muscle.

### Neonatal administration of AAV8-*Klf15* to severe SMA mice does not improve weight or survival

We next wanted to determine if increasing early postnatal *Klf15* expression in the *Smn*^*−/−*^;*SMN2* SMA mice would influence disease progression. Administering 1E11 vg/pup of AAV8-*Klf15* to P0 *Smn*^*−/−*^;*SMN2* mice and *Smn^+/−^*;*SMN2* control littermates revealed itself to be toxic (spontaneous death without any impact on weight) to both genotypes (data not shown). Seeing as this dose was not harmful with AAV8-*GFP*, the adverse effects are most likely due to the supraphysiological levels of *Klf15*. We therefore reduced the AAV8-*GFP* and AAV8-*Klf15* dose to 2E10 vg/pup for subsequent administrations, which still allowed for an age-dependent increased expression of *GFP* (Fig 2a) and *Klf15* (Fig 2b), albeit with some variability between animals, in quadriceps of P2 and P7 *Smn^+/−^*;*SMN2* and *Smn*^*−/−*^;*SMN2* mice.

**Fig 2.**
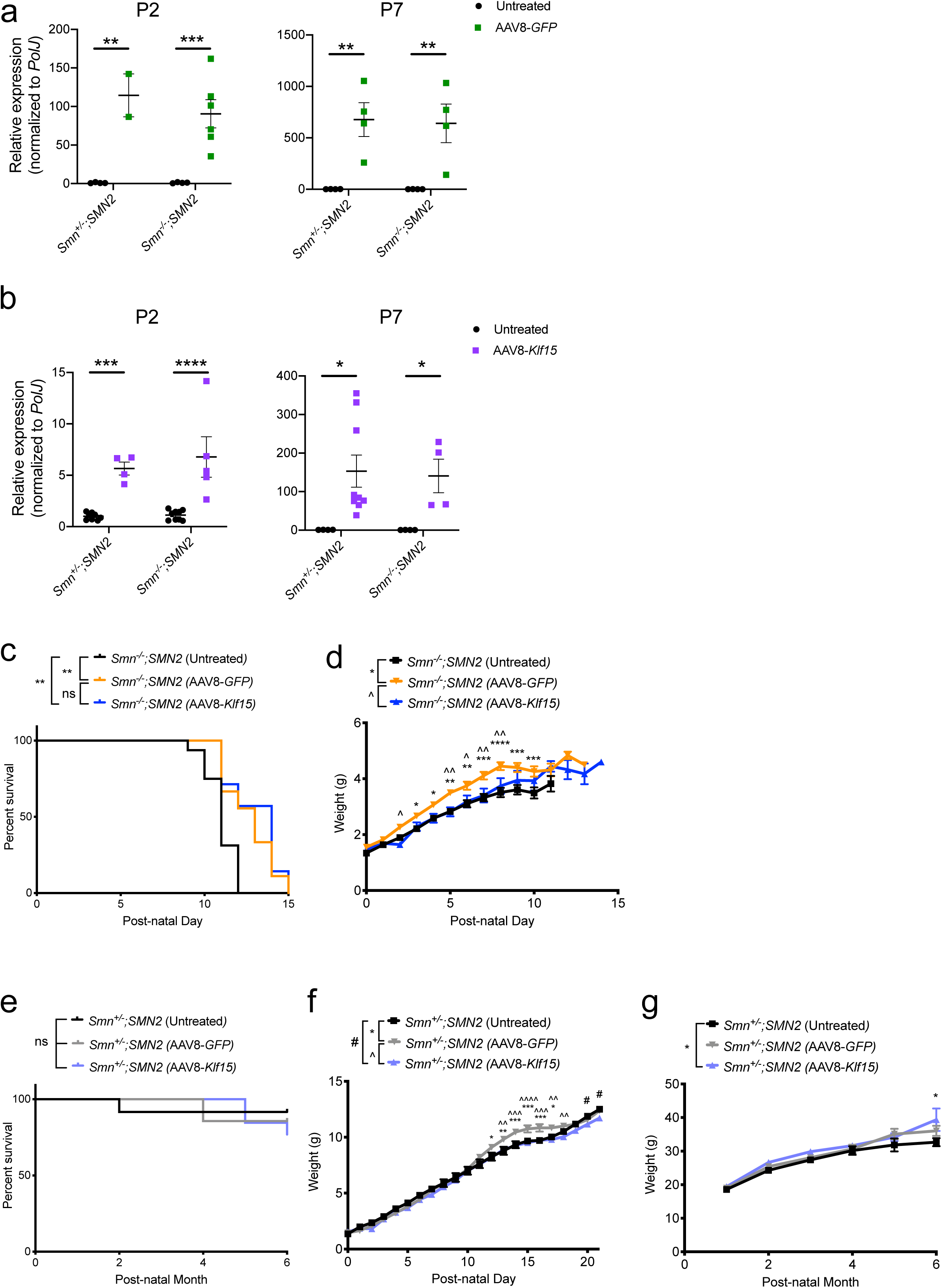
Perinatal administration of AAV8-*Klf15* does not improve weight or survival of *Smn*^*−/−*^;*SMN2* SMA mice. Post-natal day (P) 0 *Smn*^*−/−*^;*SMN2* SMA mice and control littermates were either untreated or received a single facial vein intravenous injection of AAV8-*GFP* or AAV8-*Klf15* (2E10 vg/pup). **a.** qPCR analysis of *GFP* mRNA expression in quadriceps of P2 and P7 untreated and AAV8-*GFP*-treated *Smn^+/−^*;*SMN2* and *Smn*^*−/−*^;*SMN2* mice. Data are scatter plot and mean ± SEM, n = 2–6 animals per experimental group, two-way ANOVA, \*\**p*<0.01, \*\*\**p*<0.001. **b.** qPCR analysis of *Klf15* mRNA expression in quadriceps of P2 and P7 untreated and AAV8-*Klf15*-treated *Smn^+/−^*;*SMN2* and *Smn*^*−/−*^;*SMN2* mice. Data are scatter plot and mean ± SEM, n = 4–9 animals per experimental group, two-way ANOVA, \**p*<0.05, \*\*\**p*<0.001, \*\*\*\**p*<0.0001. One outlier identified by the Grubbs’ test was removed from the P7 *Smn*^*−/−*^;*SMN2* group **c.** Survival curves of untreated (n = 16), AAV8-*GFP*-treated (n = 9) and AAV8-*Klf15*-treated (n = 7) *Smn*^*−/−*^;*SMN2* mice. Data are Kaplan-Meier survival curves, log-rank Mantel-Cox test, ns = not significant, \*\**p* = 0.0034 (untreated vs AAV8-*GFP*), \*\**p* = 0.0048 (untreated vs AAV8-*Klf15*). **d.** Daily weights of untreated (n = 16), AAV8-*GFP*-treated (n = 9) and AAV8-*Klf15*-treated (n = 7) *Smn*^*−/−*^;*SMN2* mice. Data are mean ± SEM, two-way ANOVA, */^*p*<0.05, **/^^*p*<0.01, \*\*\**p*<0.001, \*\*\*\**p*<0.0001. **e.** Survival curves of untreated (n = 12), AAV8-*GFP*-treated (n = 7) and AAV8-*Klf15*-treated (n = 13) *Smn^+/−^;SMN2* mice. Data are Kaplan-Meier survival curves, log-rank Mantel-Cox test, ns = not significant. **f.** Daily weights of untreated (n = 11), AAV8-*GFP*-treated (n = 8) and AAV8-*Klf15*-treated (n = 12) *Smn^+/−^;SMN2* mice. Data are mean ± SEM, two-way ANOVA, */^/#*p*<0.05, **/^^*p*<0.01, ***/^^^*p*<0.001, ^^^^*p*<0.0001. **g.** Monthly weights of untreated (n = 11), AAV8-*GFP*-treated (n = 7) and AAV8-*Klf15*-treated (n = 13) *Smn^+/−^;SMN2* mice. Data are mean ± SEM, two-way ANOVA, \**p*<0.05.

In terms of effects on disease progression, we found that AAV8-*Klf15*-treated *Smn*^*−/−*^;*SMN2* mice survived longer than untreated *Smn*^*−/−*^;*SMN2* (Fig 2c). However, AAV8-*GFP*-treated *Smn*^*−/−*^;*SMN2* mice also display a moderately improved lifespan (Fig 2c), suggesting that the AAV construct itself has some beneficial physiological impact. Interestingly, we also observed that *Smn*^*−/−*^;*SMN2* mice that received the AAV8-*GFP* weighed slightly more than AAV8-*Klf15*-treated and untreated *Smn*^*−/−*^;*SMN2* (Fig 2d).

We found no significant differences between the survival of untreated, AAV8-*GFP*-treated and AAV8-*Klf15*-treated *Smn^+/−^*;*SMN2* mice (Fig 2e), although a small number of spontaneous deaths occurred in all cohorts. Similar to what we observed in the *Smn*^*−/−*^;*SMN2* mice, AAV8-*GFP*-treated *Smn^+/−^*;*SMN2* mice weighed slightly more than untreated and AAV8-*Klf15*-treated *Smn^+/−^*;*SMN2* mice during the P0–P21 pre-weaning phase (Fig 2f). However, this increased weight was not maintained post-weaning (Fig 2g). We did observe a small but significant increase in weight of AAV8-*Klf15*-treated *Smn^+/−^*;*SMN2* mice at 6 months of age. Thus, administering a perinatal injection of AAV8-*Klf15* at a dose of 2E10 vg/pup significantly increases Klf15 expression in skeletal muscle without having overt adverse or beneficial effects on survival.

### Neonatal administration of AAV8-*Klf15* to intermediate SMA mice slightly improves survival

Due to the severe and rapid disease progression in the *Smn*^*−/−*^;*SMN2* mice, they respond less favorably to non-SMN treatment strategies compared to the milder intermediate *Smn*^*2B/−*^ mouse model ^18,25,32,33^. We therefore proceeded to evaluate the effect of AAV8-*Klf15* in *Smn*^*2B/−*^ mice and *Smn^2B/+^* control littermates, following the same dosing regimen as in the severe SMA mice. Similar to what was observed in the *Smn*^*−/−*^;*SMN2* mice, AAV8-*GFP*-treated *Smn*^*2B/−*^ mice also demonstrated a small but significant increase in survival compared to untreated *Smn*^*2B/−*^mice (Fig 3a). Interestingly, AAV8-*Klf15*-treated *Smn*^*2B/−*^ mice had a significantly increased lifespan compared to both untreated and AAV8-*GFP*-treated *Smn*^*2B/−*^ mice (Fig 3a). While AAV8-*Klf15* did not influence the weight of *Smn*^*2B/−*^ mice during the nursing period (up to P21), an increase in weight could be seen in the post-weaned mice (Fig 3b). However, that weight gain did not ultimately prevent an early death. Interestingly, AAV8-*GFP*-treated *Smn*^*2B/−*^ mice were significantly heavier than untreated and AAV8-*Klf15*-treated *Smn*^*2B/−*^ mice (Fig 3b) between P13–21, again similar to what we found in the *Smn*^*−/−*^;*SMN2* mice. We did not observe any effects of AAV8-*GFP* or AAV8-*Klf15* on the weights of pre-weaned *Smn*^*2B/−*^ mice compared to untreated animals (Fig 3c). Thus, the AAV8-*Klf15* was slightly more beneficial in the intermediate *Smn*^*2B/−*^ SMA mouse model than the severe Taiwanese *Smn*^*−/−*^;*SMN2* mice, overall.

**Fig 3.**
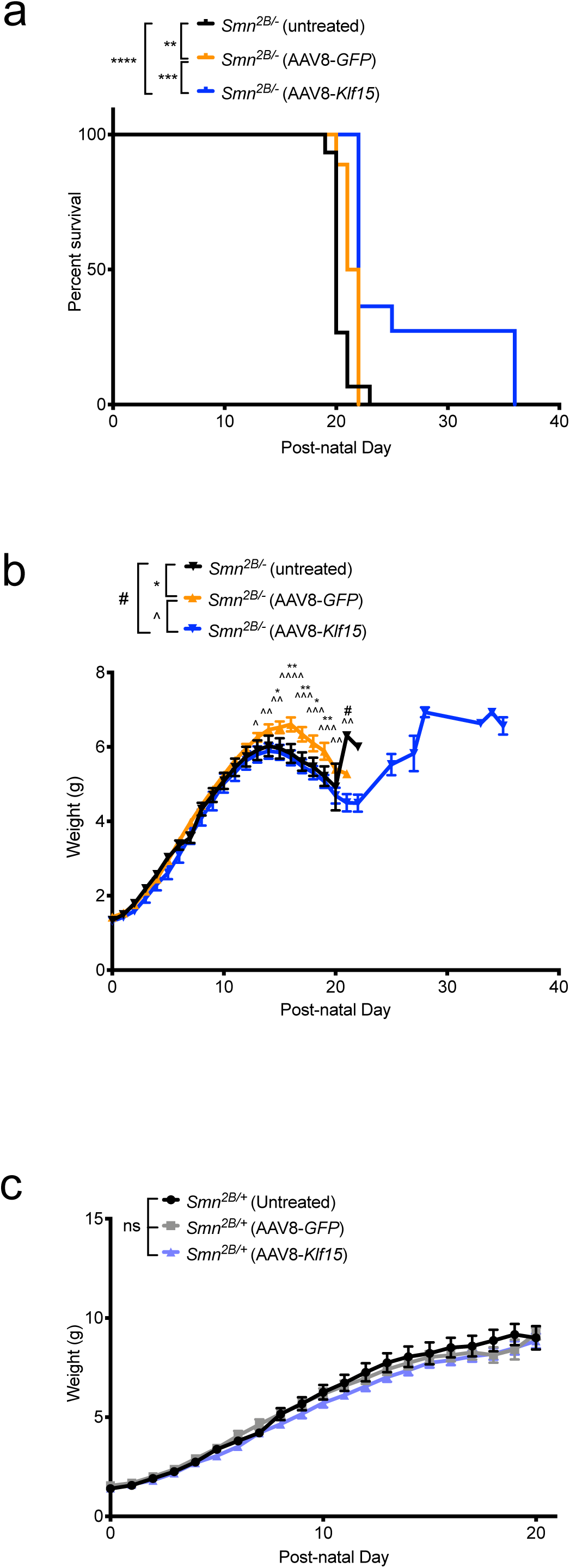
Perinatal administration of AAV8-*Klf15* slightly increases survival in *Smn*^*2B/−*^ SMA mice. Post-natal day (P) 0 *Smn*^*2B/−*^ SMA mice and control littermates were either untreated or received a single facial vein intravenous injection of AAV8-*GFP* or AAV8-*Klf15* (2E10 vg/pup). **a.** Survival curves of untreated (n = 15), AAV8-*GFP*-treated (n = 18) and AAV8-*Klf15*-treated (n = 11) *Smn*^*2B/−*^ mice. Data are Kaplan-Meier survival curves, log-rank Mantel-Cox test, \*\**p* = 0.0021 (untreated vs AAV8-*GFP*), \*\*\*\**p*<0.0001 (untreated vs AAV8-*Klf15*), \*\*\**p* = 0.0008 (AAV8-*GFP* vs AAV8-*Klf15*). **b.** Daily weights of untreated (n = 18), AAV8-*GFP*-treated (n = 18) and AAV8-*Klf15*-treated (n = 11) *Smn*^*2B/−*^ mice. Data are mean ± SEM, two-way ANOVA, */^/#*p*<0.05, **/^^*p*<0.01, ^^^*p*<0.001, ^^^^*p*<0.0001. **c.** Daily weights of untreated (n = 11), AAV8-*GFP*-treated (n = 9) and AAV8-*Klf15*-treated (n = 16) *Smn^2B/+^* mice. Data are mean ± SEM, two-way ANOVA, ns = not significant.

### The AAV8-Spc5-12 construct also induces expression in heart and liver

While the synthetic Spc5-12 promoter has been used for its enhanced activity in skeletal muscle ^27,31^, we wanted to determine if our AAV8 delivery system also induced expression in heart and liver, tissues in which *Klf15* also plays key roles ^34,35^. We indeed observed significant increased expression of *GFP* and *Klf15* mRNA in the livers of both P2 and P7 AAV8-*GFP*-(Fig 4a) and AAV8-*Klf15*-treated (Fig 4b) *Smn^+/−^*;*SMN2* and *Smn*^*−/−*^;*SMN2* mice compared to untreated animals. This increase was approximately 15 times (*GFP*) and 10 times (*Klf15*) higher than in skeletal muscle for both time points. Similarly, we find a significant upregulation of *GFP* and *Klf15* mRNA in the hearts of AAV9-*GFP*-(Fig 4c) and AAV8-*Klf15*-treated (Fig 4d) P2 and P7 *Smn^+/−^*;*SMN2* and *Smn*^*−/−*^;*SMN2* mice compared to untreated animals. Surprisingly, the increase in heart was approximately 40 times (*GFP*) and 50 times (*Klf15*) higher at P2 while in was approximately 70 times (*GFP*) and 10 times (*Klf15*) higher at P7 compared to skeletal muscle at those respective time points. Of note, there was some variability between animals within the same experimental group. Therefore, our results demonstrate that the activity of the AAV8-Spc5-12 vector is not exclusive to skeletal muscle.

**Fig 4.**
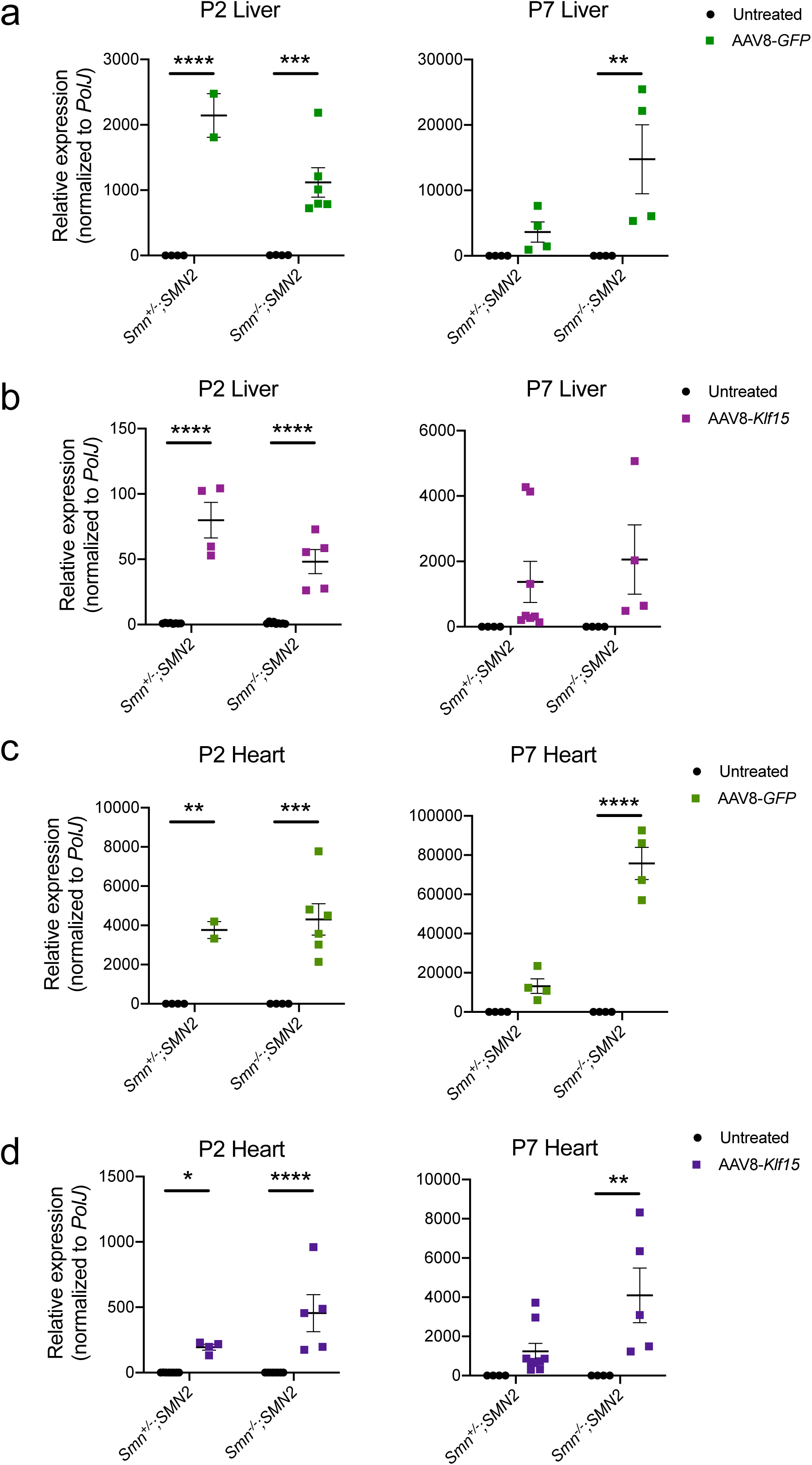
Perinatal administration of the AAV8-Spc5-12 construct induces high expression in liver and heart. Post-natal day (P) 0 *Smn*^*−/−*^;*SMN2* SMA mice and control littermates were either untreated or received a single facial vein intravenous injection of AAV8-*GFP* or AAV8-*Klf15* (2E10 vg/pup). **a-b.** qPCR analysis of *GFP* (**a**) and *Klf15* (**b**) mRNA expression in quadriceps muscles of P2 and P7 untreated and AAV8-treated *Smn^+/−^*;*SMN2* and *Smn*^*−/−*^;*SMN2* mice. Data are scatter plot and mean ± SEM, n = 2–6 animals per experimental group, two-way ANOVA, \*\**p*<0.01, \*\*\**p*<0.001, \*\*\*\**p*<0.0001. One outlier identified by the Grubbs’ test was removed from the P7 liver AAV8-*Klf15 Smn^−/−^;SMN2* group. **c-d.** qPCR analysis of *GFP* (**c**) and *Klf15* (**d**) mRNA expression in heart of P2 and P7 untreated and AAV8-treated *Smn^+/−^*;*SMN2* and *Smn*^*−/−*^;*SMN2* mice. Data are scatter plot and mean ± SEM, n = 2–8 animals per experimental group, two-way ANOVA, \*\**p*<0.01, \*\**p*<0.01, \*\*\**p*<0.001 \*\*\*\**p*<0.0001. One outlier identified by the Grubbs’ test was removed from the P2 heart untreated *Smn*^*−/−*^;*SMN2* group.

## DISCUSSION

We have recently demonstrated that *Klf15* expression is significantly downregulated in muscle of pre-symptomatic SMA mice and that upregulating *Klf15* expression via genetic (transgenic muscle-specific expression) or pharmacological (prednisolone) approaches results in improved disease phenotypes ^18^. Here, we evaluated the impact of specifically upregulating *Klf15* in skeletal muscle in perinatal mice by driving its expression via an AAV8-Spc5-12 vector. We find that while neonatal administration of the AAV8-*Klf15* construct leads to significant increased levels of *Klf15* in muscle, this has no overt effect on survival or weight gain in the severe Taiwanese SMA mice while we observe a small improvement in the lifespan of the intermediate *Smn*^*2B/−*^ mice.

The lack of significant impact of AAV8-*Klf15* in severe SMA mice may be due to several compounding factors. While the levels of *Klf15* expression achieved with AAV8-*Klf15* are similar to the levels observed in transgenic SMA mice overexpressing muscle-specific *Klf15* at P2 (~ 5–10 fold greater than control littermates), the amounts measured in P7 AAV8-*Klf15*-treated animals are significantly greater than the transgenic mice (~ 65–740 fold greater and ~ 30 fold greater, respectively, compared to control littermates) (Fig 2) ^18^. As *Klf15* can display both atrophy-inducing ^36^ and ergogenic ^19^ properties in skeletal muscle in a dose-dependent manner ^37^, it is quite possible that the supraphysiological levels achieved with AAV8-*Klf15* favor muscle wasting over growth. We also note significantly more variability in *Klf15* levels in animals injected with the AAV8 construct compared to the transgenic mice (Fig 2) ^18^, which are most likely due to differential injection efficiencies and/or vector spread and could influence physiological outcomes.

We have previously shown that administration of prednisolone to SMA mice also increases *Klf15* levels in skeletal muscle of P2 pre-symptomatic animals (~ 6 fold greater than untreated controls) ^18^. However, this effect on *Klf15* induction ceased at P7, specifically in SMA mice ^18^, suggesting that prednisolone-dependent benefits in symptomatic SMA mice may be due to KLF15-independent effects and/or that prednisolone-dependent *Klf15* increase in P7 animals may be limited by compensatory inhibitory mechanisms due to already significantly increased *Klf15* levels in symptomatic SMA mice compared to controls ^18^. It is therefore possible that an optimal strategy would be to conditionally increase *Klf15* expression in pre-symptomatic stages only, which is not easily achieved as the kinetics of AAV-mediated overexpression require several days for efficient transgene activity. To achieve optimal expression at early pre-symptomatic post-natal stages may therefore require pre-natal delivery.

While our AAV8 construct was designed to overexpress *Klf15* specifically in skeletal muscle, our analysis of heart and liver demonstrate significantly higher activity in these tissues (Fig 4). KLF15 has well described roles in heart and liver ^34,35^, suggesting that its increased expression via our AAV8 construct, most likely also impacts the function of these tissues. SMA mice display several heart and liver pathologies ^38,39^, suggesting that aberrant KLF15 levels may have non-intended organ-specific adverse effects, which most likely explain the spontaneous deaths observed in our mice treated with the high vector dose. Furthermore, we have previously reported increased levels of *Klf15* in the liver and heart of symptomatic mice ^18^, which most likely reflect a pathological response.

Surprisingly, the AAV8-*GFP* construct also demonstrated some non-negligible effects on disease phenotypes of SMA mice (Fig 3). While we cannot be certain as to why that is, we speculate that it may be related to possible effects on the immune system, similarly to previous reports for AAV8 vectors ^40,41^. Seeing as SMA mice have an altered immune response ^42,43^ and that inflammation can display both protective and adverse systemic properties ^44^, including the CNS ^45^ and muscle ^46^, it is possible that an activated immune response results in acute and/or intermittent benefits in AAV8-treated SMA animals.

In summary, the limited impact of AAV8-*Klf15* administration in SMA mice might be explained by several experimental conditions that most likely reduced our ability to increase Klf15 specifically in skeletal muscle at physiological levels and with the optimal timing, without influencing the function of other tissues and systems. In the experimental paradigms tested here, the positive, albeit small, effect on survival and weight was restricted to the milder *Smn*^*2B/−*^ SMA mouse model.

## ACKNOWLEDGMENTS

NA, DH, RJYM and MB were supported by the UK SMA Research Consortium SMA Trust UK grant. SK was supported by an ERASMUS grant and PC received financial support from the Deutsche Muskelstiftung.

